# Pulsed ELDOR Measurement of the Distance Between a Spin-Label and Copper (II) Centre in the Copper Loaded R48C Mutant of *N. gonorrhoeae* Ferric Binding Protein

**DOI:** 10.1101/248054

**Authors:** Matt Bawn, Justin Bradley, Fraser MacMillan

## Abstract

Distance determination in proteins and biomolecules using pulsed EPR (electron paramagnetic resonance) techniques is becoming an increasingly popular and accessible technique. PELDOR (pulsed electron-electron double resonance), is a technique designed for distance determination over a nanoscopic scale. Here, ferric binding protein (Fbp) is used to demonstrate the practicability of this technique to Cu (II) Metalloproteins. PELDOR is usually applied to bi-radicals or endogenous radicals, and distance determination using pulsed EPR of metal containing centres in biomolecules has been restricted to relaxation experiments. PELDOR distance measurements between a Cu (II) ion and a nitroxide have previously only been reported for model compounds [1, 2].

Fbp as the name suggests usually, contains a Fe (III) ion centre. For the purposes of this investigation the Fe (III) ion was removed and replaced by a Cu (II) ion, after a nitroxide spin-label was added to the Fbp using of site directed spin-labelling (SDSL). PELDOR was then applied to measure the distance between the two centres.

Simulation methods were then employed to fully investigate these data and allow a quantitative interpretation of the copper nitroxide PELDOR data. The observed PELDOR time traces were analysed using DEER analysis[3].

## Results

Ferric binding proteins (FBPs) are efficient iron sequestering proteins produced by many pathogenic bacteria such as *Neissaria gonorrheae^1^* to procure iron from their hosts. FBPs have considerable structural similarity to one lobe of transferrin,^2^ but without sequence homology.^3^ The Fe (III) binding site in *holo*-FBP (hFBP) has been found to be similar to the two Fe (III) binding sites in transferrin, each containing two *cis*-Tyr and one His ligand, but with FBP containing a Glu residue instead of the Asp found in transferrin.^4^

Pulsed electron double resonance (PELDOR) is a technique that is able to monitor the dipole interaction, v_dd_, between two paramagnetic centers.^5,6^ This interaction may then be used to determine the distance between the two centers as shown by the equation:

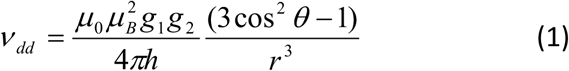

Where g_1_ and g_2_ are the g-values of the two paramagnetic centers, r is the distance between the two centers and *ϑ* is the angle between the external magnetic field and the spin-spin vector. This technique may be used to determine distances in the range 15 to 80 A^°^.^7^

PELDOR was initially, and still is^8^ predominantly used to measure the distance between two nitroxide spin labels, however, as the technique has developed it has also been applied to proteins containing metal centers. This is generally experimentally more challenging due to the relaxation properties of the metal ion, and the anisotropy of its g-tensor. For this reason the determination of the distance between a Cu (II) center and a spin-label has previously been performed only on model systems.^9,10^ This communication describes the first example of the application to a biological macromolecule.

Figure 1 displays the X-Band (9 GHz) field-swept electron-spin echo spectrum of (a) Cu (II) loaded spin-labeled Fbp R48C, (b) the simulated spectrum for this frequency using the EasySpin^11^ toolbox for Matlab, and (c) the contributions to the simulation of the Cu (II) center (red) and the spin-label (blue). From this it can clearly be seen that the position of the pump pulse corresponds to the maximum intensity of the spin-label contribution, whilst the Cu (II) center was chosen as the detection location. To minimize the effects of orientation selectivity the g⊥ position on the Cu (II) spectrum was chosen^12^. At this position the contribution to the spectrum from a wide range of orientations is superimposed. Δ*v* (the difference in frequency between the pump and detect pulses) was 132 MHz (the experimental constraint on the choice of this value is the operational bandwidth of the resonator).

**Figure 1.**
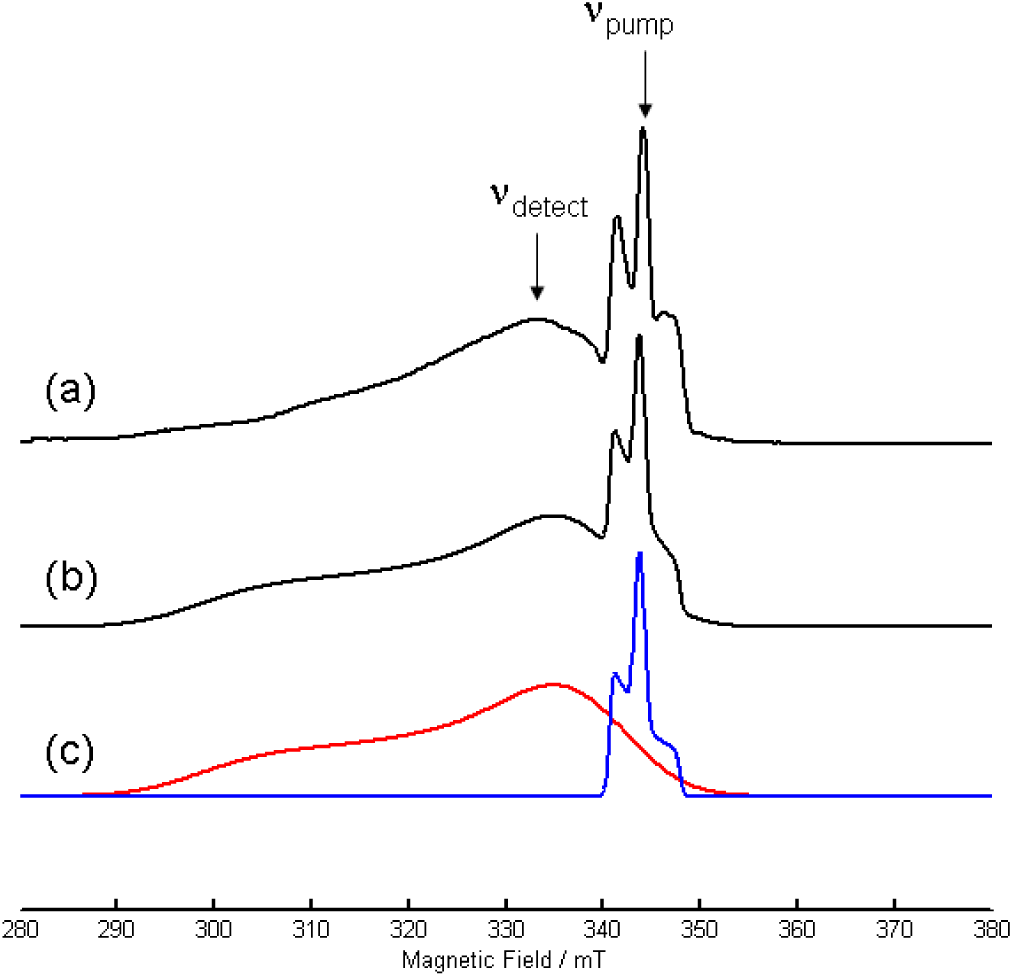
(a) Field-swept electron-spin echo spectrum of Cu (II) Fbp R48C recorded at 10 K with an X-band Bruker ELEXSYS E680 spectrometer operating at 9.672 GHz.. A two-pulse echo sequence (π/2-τ-π) with a 16 ns π-pulse and τ = 200 ns was used. Arrows represent positions of pumping and detection. (b) A Matlab simulation of Cu (II) Fbp R38C using the EasySpin toolbox. (c) The contributions to (b) from the Cu (II) paramagnetic center (red) and the nitroxide spin-label (blue).

To determine the effect of orientation selectivity to the generated distance distribution, the PELDOR experiment was repeated with Δv = 500 MHz. Figure 2. shows the time domain PELDOR spectra of Cu (II) Fbp R48C measured at 10 K, with (a) Δ*v* = 132 MHz, and (b) Δ*v* = 500 MHz. With a larger offset frequency the number of Cu (II) spins excited by the detect pulse was reduced, leading to a higher degree of noise in spectrum 2 (b).

**Figure 2.**
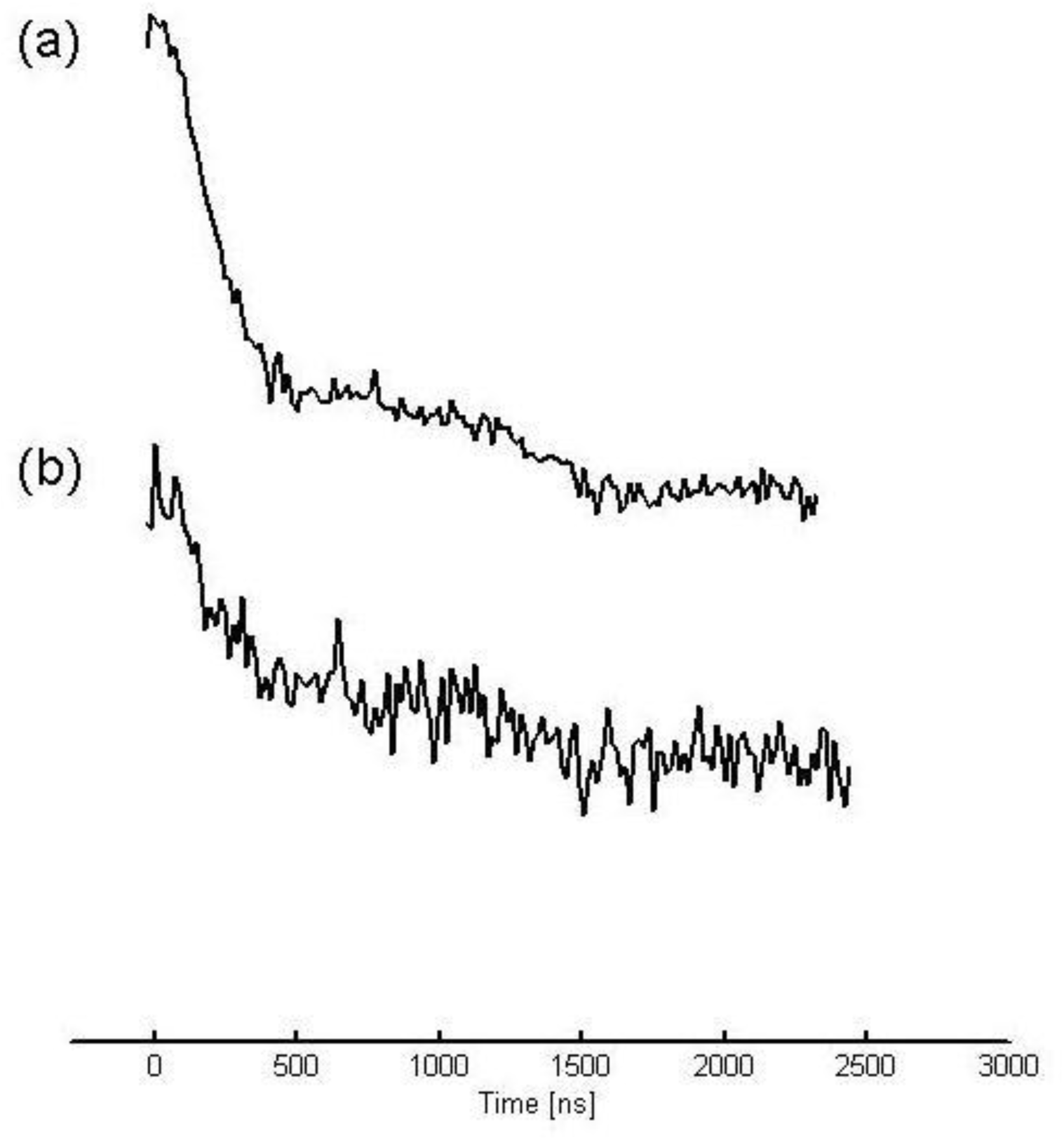
Four-Pulse PELDOR spectrum of Cu (II) Fbp R48C recorded at 10 K with X-band Bruker ELEXSYS E680 spectrometer with a 32 ns π-pulse. (a) Δ*V* = 132 MHz. (b) Δ*V* = 500MHz.

DeerAnalysis 2009^13^ was then used to analyze the time domain spectra. This program, also based in Matlab, allows distance distributions to be extracted from dead-time free PELDOR data. Tikhonov regularization with a regularization parameter of 100 on the L-curve was used to generate the distance distributions. These can be seen in Figure 3. with the black trace showing the distributions from the PELDOR data with Δv = 132 MHz, and the red trace showing that from the data with Δ*v* = 500 MHz. It can clearly be seen that doth distributions show markedly similar characteristics. The plot also indicates the position of the mean distance of 3.65 nm taken from the Δ*v* = 132 MHz trace. The extra spectral features on the red curve are due to artifacts originating from Tikhonov regularization of significantly noisier data.

**Figure 3.**
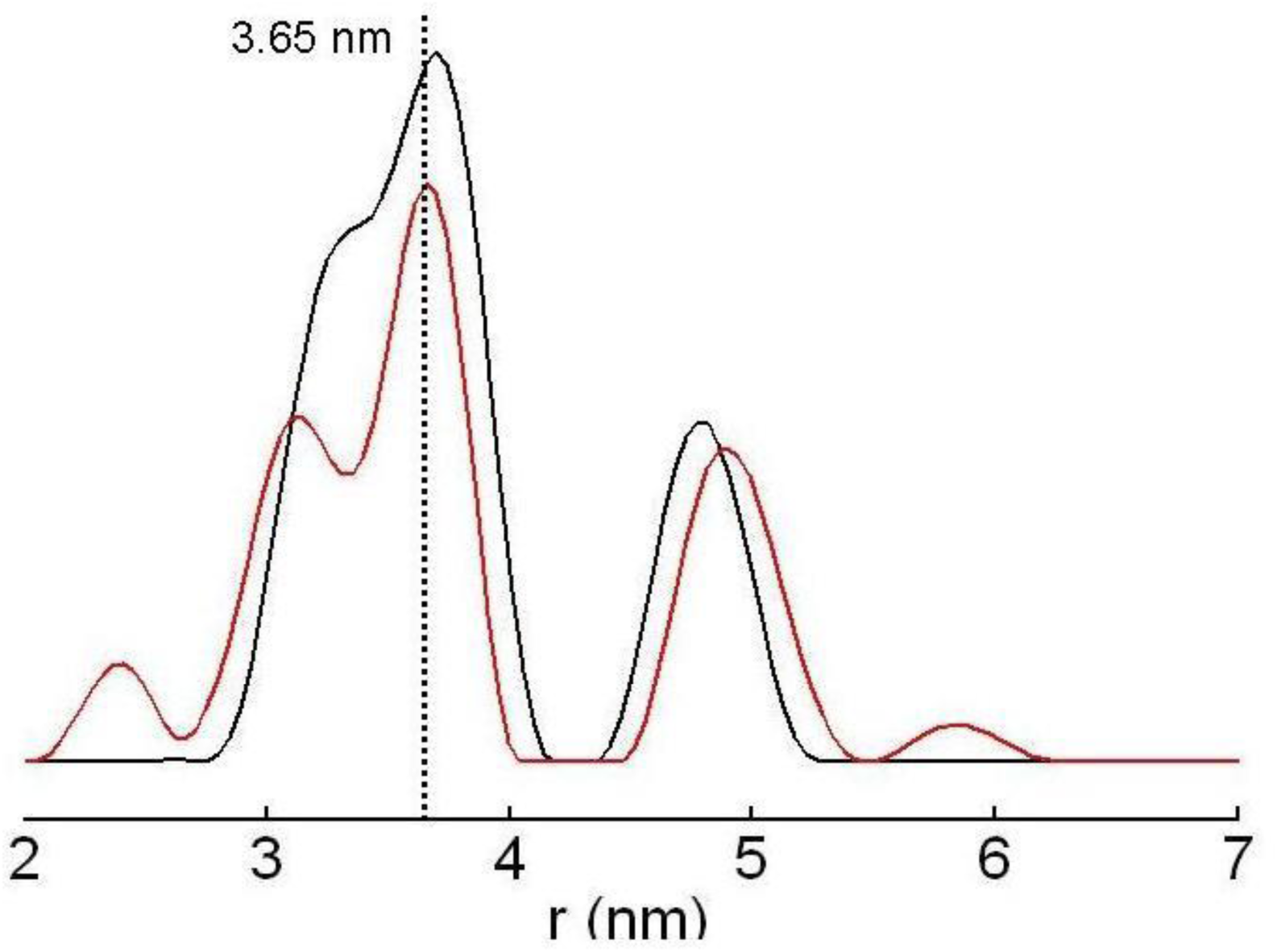
Distance distributions derived from Tikhonov regularization, with regularization parameter 100. Black curve indicates distribution from experiment performed with Δ*V* = 135 MHz, red curve with Δ*V* = 500 MHz.

An examination of the crystal structure of FbpA from *N. gonorrheae* (PDB 1d9y unpublished) yields a metal center to R48 residue distance of 2.8 nm, and the length of the spin-label used in this study was 0.7 nm. Addition of these two distances produces a theoretical Cu (II) to spin label distance of 3.5 nm. This compares well to the mean distance determined by PELDOR of 3.65 nm.

In conclusion this communication shows that PELDOR may be used to determine distance distributions between Cu (II) centers and spin-labels in a biologically relevant protein.

## ACKNOWLEDGMENT

Dominic Campopiano is acknowledged for providing the Fbp R48C mutant and Dr Gaye White for her help and guidance. The EPRSC is also recognized for the provision of a studentship.

